# Transcription factor TFAP2C affects porcine early embryo development via regulating epigenetic modification

**DOI:** 10.1101/2022.11.25.517906

**Authors:** Daoyu Zhang, Di Wu, Sheng Zhang, Meng Zhang, Yongfeng Zhou, Xinglan An, Qi Li, Ziyi Li

**Affiliations:** Key Laboratory of Organ Regeneration and Transplantation of Ministry of Education, First Hospital, Jilin University, Changchun 130021, China; First Hospital, Jilin University, Changchun 130021, China; The Jackson Laboratory for Genome Technology, 10 Discovery Drive Farmington, Connecticut, 06932, USA

**Keywords:** TFAP2C, porcine embryo development, histone modification, *SETD2*

## Abstract

Transcription factors (TFs) have the potential function in regulating gene expression. Transcription factor TFAP2C plays important roles in the regulation of post-implantation embryonic development in mice, the reprogramming process, trophectoderm formation and carcinogenesis, but its role in porcine early embryo development remains unclear. This study was conducted to investigate the role of TFAP2C in porcine early embryo development using siRNA cytoplasmic injection. The RNAseq and immunofluorescence staining were performed to detect gene expression, and ChIP and dual luciferase reporter assays were used to elucidate the mechanism. The results showed that the deficiency of TFAP2C could lead to embryonic development disorder. The percentage of the blastocyst in the *TFAP2C* knockdown (*TFAP2C-KD*) group (7.76±1.86%) was significantly decreased compared to the control group (22.92±1.97%) (*P*\*\*<0.01). The RNAseq results showed that 1208 genes were downregulated and 792 genes were upregulated after siRNA injection. The expression of epigenetic modification enzymes KDM5B, SETD2 (*P*\*\*<0.01) *etc*. were significantly elevated in *TFAP2C-KD* group. Meanwhile, the modification levels of H3K4me3, H3K4me2 and H3K9me3 (*P*\*<0.05) were significantly decreased, and the modification levels of H3K36me3 (*P*\*\*<0.01) and DNA methylation (*P*\*\*<0.01) were significantly increased in *TFAP2C*-KD group. DNMT1 was mostly expressed in cytoplasm in the control group, while it was mainly expressed in nuclei in the *TFAP2C*-KD group. In addition, TFAP2C could bind to the promoter region of *SETD2*, and the mutation of the TFAP2C binding site resulted in increased activity of *SETD2* promoter (*P*\*\*<0.01). The knockdown of TFAP2C affects histone modification and DNA methylation by regulating the expression of *SETD2, KDM5B etc*. and other genes, thereby inhibiting embryonic development. TFAP2C binds to the promoter region of *SETD2* and acts as a hindrance protein. This study fills in the deficiency of TFAP2C in porcine early embryo development and provides theoretical support for animal husbandry production and biomedicine.

**Author Summary:** The correct activation of embryonic genes is required during early embryonic development, and the activation of these genes is subject to strict epigenetic regulation, such as DNA methylation, histone acetylation and methylation, with abnormalities in either leading to birth defects and developmental defects in individuals. TFs have specific binding motifs that regulate gene expression by binding to them. TFAP2C has been studied in post-implantation embryonic development and trophectoderm generation, however, the effect on early embryo development is unknown. Our findings suggest that TFAP2C deficiency disrupts gene expression patterns and leads to abnormal epigenetic modifications, resulting in abnormal embryo development. Furthermore, we found for the first time that TFAP2C can bind to the promoter region of *SETD2*, thereby affecting early embryo development in pigs. This indicates the critical role of TFAP2C in early embryo development in pigs on one hand, and also provides theoretical support for livestock production and biomedicine.

## Introduction

From the beginning of fertilization to the end of implantation, embryonic development can be divided into several carefully arranged stages: fertilization, mulberry embryo and blastocyst formation. Understanding the stages of early embryo development and the underlying molecular mechanisms of regulation is essential for basic reproductive biology and practical applications including regenerative medicine. It is essential to determine the early embryo’s overall gene, RNA and protein expression patterns for deciphering the regulatory mechanisms (1). Transcription factors (TFs) control chromatin and transcription by recognizing specific DNA sequences, despite the strong interest of many scientists in understanding how transcription factors control gene expression, however, the role of transcription factors in early embryo development still needs a lot of research to explore.

TFAP2C, a member of the AP-2 family, plays a role in several aspects of tumorigenesis, stem cell maintenance and embryonic development (2). It has a canonical binding motif GCCNNNGGC. Previous studies have shown that *TFAP2C* is significantly induced during trophoblastic ectodermal differentiation in humans and murine (3, 4). Winger *et al*. reports that *TFAP2C* is expressed during mouse preimplantation embryonic development, is expressed in oocytes and declines sharply after fertilization, resumed expression at the 4-cell stage and persists through the blastocyst stage (5). Moreover, *TFAP2C* is required for the survival of the mouse embryo, *TFAP2C* deficiency leads to mouse embryonic lethality at approximately embryonic day (E)7.5, which may be attributed to defective placental development (6, 7). However, studies of TFAP2C in early embryo development are scarce.

Histone modifications are associated with the activation and repression of transcription. Earlier studies report their functions in the regulation of transcriptional, for example, H3K4me1(8) and H3K4me3 (9) are associated with transcriptional activation, while H3K9me3 and H3K27me3 are associated with transcriptional repression (10). H3K4me3 is localized in the promoter region of the gene and is centered at the transcriptional start site (11). In mouse embryonic stem cells (mESCs), most transcription start sites are found to be tagged by H3K4me3. In the same study, they find that H3K27me3 is more widely distributed, but that 75% of the H3K27me3 sites spanning the transcription start site (TSS) are also labeled by H3K4me3 (12). In embryonic development, H3K4me3 leads to gene transcription from developing gametes to post-implantation embryos (13). A previous study reports that the downregulation of H3K4me3 in full-grown oocytes by overexpression of the H3K4me3 demethylase KDM5B is associated with defects in genome silencing (13).

Cytosine methylation in mammalian cells occurs mainly in CpG dinucleotides, and molecular and genetic studies have shown that DNA cytosine methylation (abbreviated as 5mC for 5-methylcytosine) is associated with gene silencing (14). DNA methylation is mainly established and maintained by DNA methyltransferases (DNMTs), that transfer a methyl group from S-adenyl methionine (SAM) to the fifth carbon of a cytosine residue to form 5mC (15, 16). While the active removal of methylation marks relies on the activity of ten-eleven translocation enzymes (TET) and thymine DNA glycosylase (TDG) (17, 18). DNMT1s are associated with MII oocyte chromatin and are present in the nucleus throughout preimplantation development. Recent studies indicate that methylation of CpG islands relates to the methylation status of H3K4 (11), and the levels of methylated H3K4 (H3K4me3) tend to be inversely correlated with DNA methylation (19).

Pigs are important livestock for agriculture and a valuable animal model for xenotransplantation, disease models and basic biology research. After fertilization, normal chromatin reprogramming is essential for embryonic development, and it is commonly believed that embryonic genome activation occurs at the 4-cell stage of the porcine embryos. In this study, the key TFs that regulate embryo development were screened by analyzing the expression pattern of TFs in porcine embryos. And TFAP2C was screened for follow-up studies, by knocking down *TFAP2C* in embryos, we investigated its function and mechanism of regulating embryonic development. This study fills in the deficiency of TFAP2C in porcine early embryo development and provides theoretical support for biomedicine and animal husbandry production.

## Results

### 2.1 TFAP2C was highly expressed from 4-cell to blastocyst in porcine embryos

To search for key TFs during embryonic development, we analyzed the RNAseq data of MII oocytes and IVV embryos including 2-cell, 4-cell, 8-cell, blastocyst. And as shown in figure 1A, the heatmap showed the expression pattern of transcription factors in IVV embryos. There was a total of 23 genes, among which 9 genes were highly expressed in oocytes and the 2-cell stage, while they were lowly expressed in 4-cell, 8-cell and blastocyst stages. There were 12 genes lowly expressed in oocytes and the 2-cell stage, while they were highly expressed in 4-cell, 8-cell and blastocyst stages. We focused on TFAP2C from these TFs, firstly, it was upregulated from the 4-cell stage (figure 1B). In addition, the function of TFAP2C in embryonic development remained unclear. Next, we designed two siRNA of *TFAP2C*, and transfected pig fibroblasts with them and assayed their interference efficiency. As shown in figure 1C, the *TFAP2C*-01 had a better interference effect.

**Figure 1.**
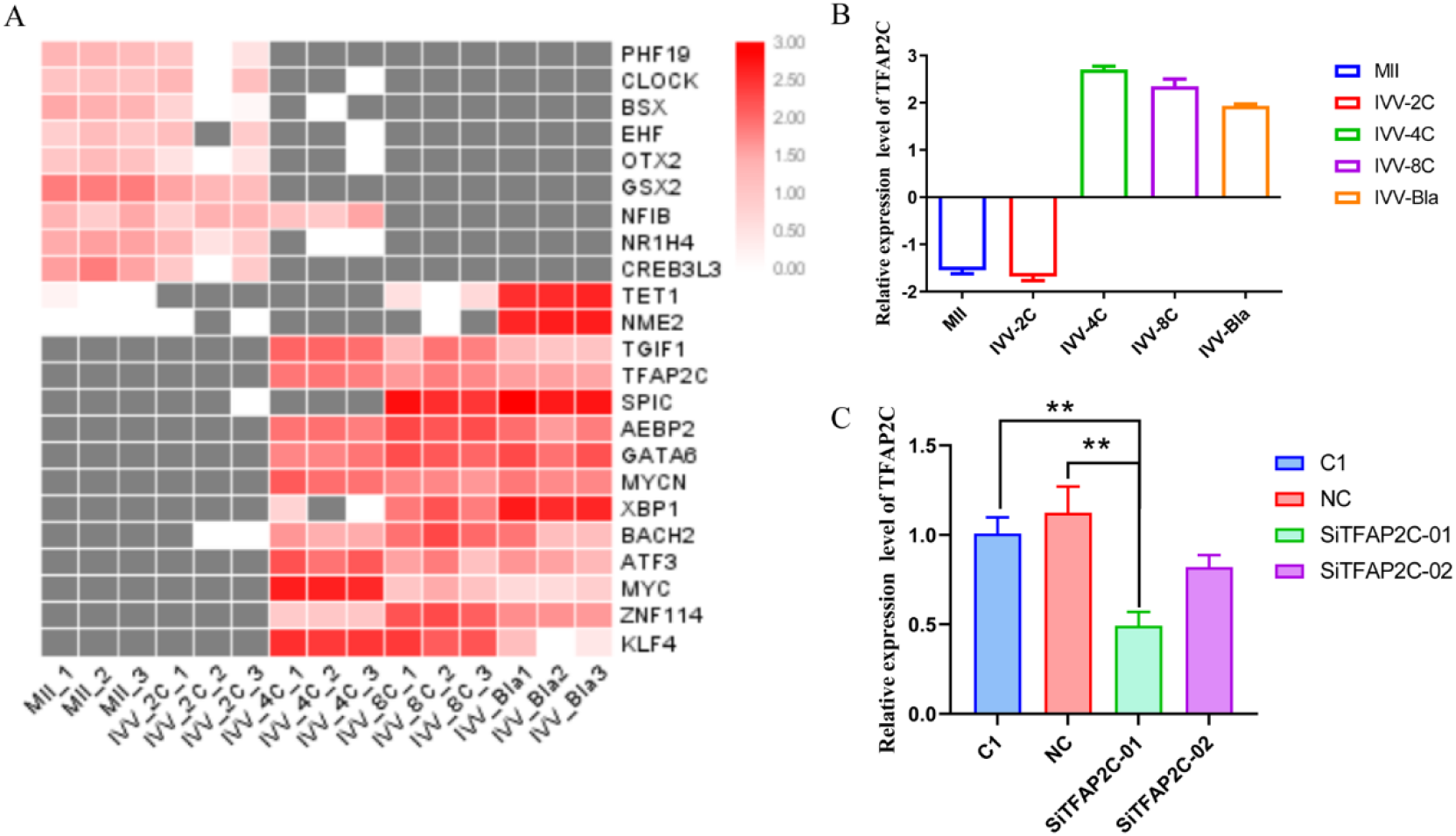
High expressed level of TFAP2C in porcine embryos from 4-cell to blastocyst. (A) The heatmap of transcription factors in pig IVV embryos. (B) The expression of *TFAP2C* in MII oocytes, and 2-cell, 4-cell, 8-cell and blastocyst in IVV embryos. (C) Detection of interference effect of siRNAs.

### 2.2 The knockdown of TFAP2C inhibited porcine embryonic development

To investigate the function of TFAP2C in embryo development, the siRNA of *TFAP2C* were injected into the MII oocytes, which were then fertilized in vitro (figure 2A), and its effect on the embryonic development were investigated. As shown in figure 2B and Table 1, no difference was observed in the percentage of the blastocyst between NC injected group (20.4±0.95%) and the control group (22.92±1.97%), suggesting there was no significant effect of NC. But the percentage of the blastocyst in the *TFAP2C* knockdown (*TFAP2C-KD*) group (7.76±1.86%) was significantly decreased compared to the control group (*P*\*\*<0.01). Meanwhile, we also observed that the embryonic development difference was significantly changed from the 8-cell stage (figure 2C) (*P*\*\*<0.01). As expected, the TFAP2C was decreased in *TFAP2C-KD* embryos compared to NC or control (figure 2D–2E) (*P*\*\*\*<0.001). We also observed more apoptosis cells in *TFAP2C*-KD embryos (figure 2F). These data indicated that the knockdown of *TFAP2C* inhibited porcine early embryo development.

**Figure 2.**
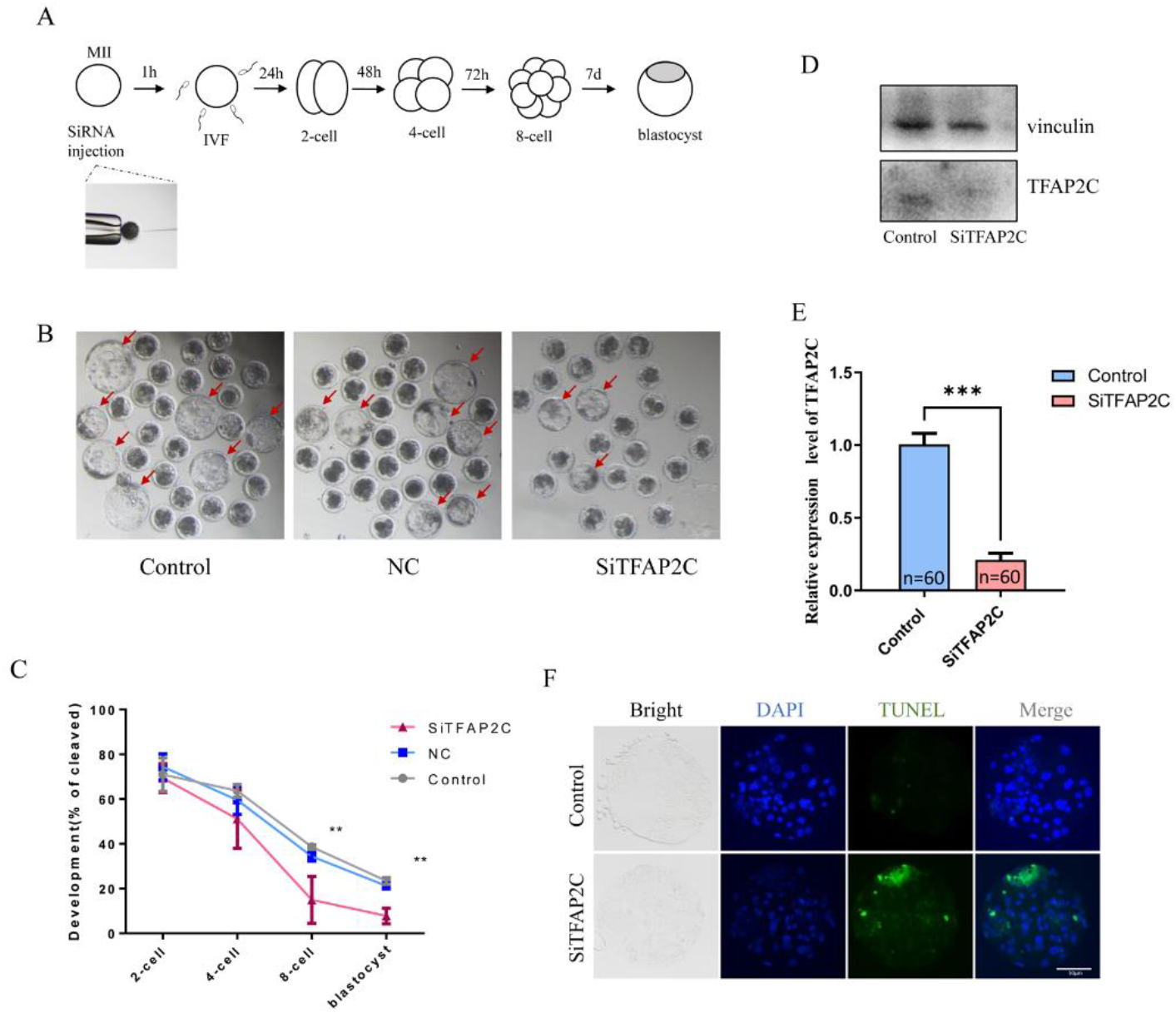
Interference with TFAP2C inhibited IVF embryo development. (A) Flow chart of interfering RNA injection and observation of early embryo development. (B) The three images were the blastocyst of control group, NC group and TFAP2C-KD group, respectively. (C) The embryonic development rate in 2-cell, 4-cell, 8-cell and blastocyst, *P*\*\*<0.01. (D) Western blotting analysis of TFAP2C expression of control group and TFAP2C-KD group. (E) qPCR analysis of *TFAP2C* expression of control and *TFAP2C-KD* groups at 48 h after siRNA injection (2-cell stage, n=60 per group). The experiment was independently repeated three times. (F) The TUNEL stain of control group and *TFAP2C-KD* group. The green indicated apoptosis cells. The experiment was repeated three times independently with 6-10 embryos/stage /group /replicate.

**Table 1:** The development of porcine IVF embryos in control, NC and *Si-TFAP2C* groups. The developmental rates of control group and NC group were also showed in our other article (42).

|  | Experiments | Embryos | 2-cell | 4-cell | 8-cell | Blastocyst |
| --- | --- | --- | --- | --- | --- | --- |
|  | (n) | (n) | (Mean±SEM%) | (Mean±SEM%) | (Mean±SEM%) | (Mean±SEM%) |
| Control | 4 | 445 | 70.95±3.25 | 63.7±1.24 | 35.59±2.00<sup>a</sup> | 22.92±1.97<sup>a</sup> |
| NC | 3 | 283 | 74.42±2.50 | 59.5±2.71 | 34.35±0.96<sup>a</sup> | 20.4±0.95<sup>a</sup> |
| Si-TFAP2C | 3 | 270 | 69.36±2.77 | 51.15±5.67 | 14.93±4.9<sup>b</sup> | 7.76±1.86<sup>b</sup> |

a, b Values with different superscript letters within a column differ significantly (P<0.05). The experiment was replicated at least three times.

### 2.3 The knockdown of TFAP2C disturbed transcriptional reprogramming in embryos

In order to explore the regulating mechanism of TFAP2C in porcine embryonic development, the 4-8 cell stage of embryos were collected for RNAseq. As shown in figure 3A, the volcano showed the DEGs between the control group and the *TFAP2C-KD* group, there were 1208 genes downregulated and 792 genes upregulated in *TFAP2C-KD* embryos. We analyzed the function of these genes, as shown in figure 3B, the upregulated genes were mainly enriched in the function of regulation of intrinsic apoptotic signaling pathway, peptide biosynthetic process, translation, amide biosynthetic process *etc*.. While the downregulated genes were enriched in negative regulation of reactive oxygen species metabolic process, regulation of protein localization to cell periphery, cortical cytoskeleton organization, receptor recycling *etc*..

**Figure 3.**
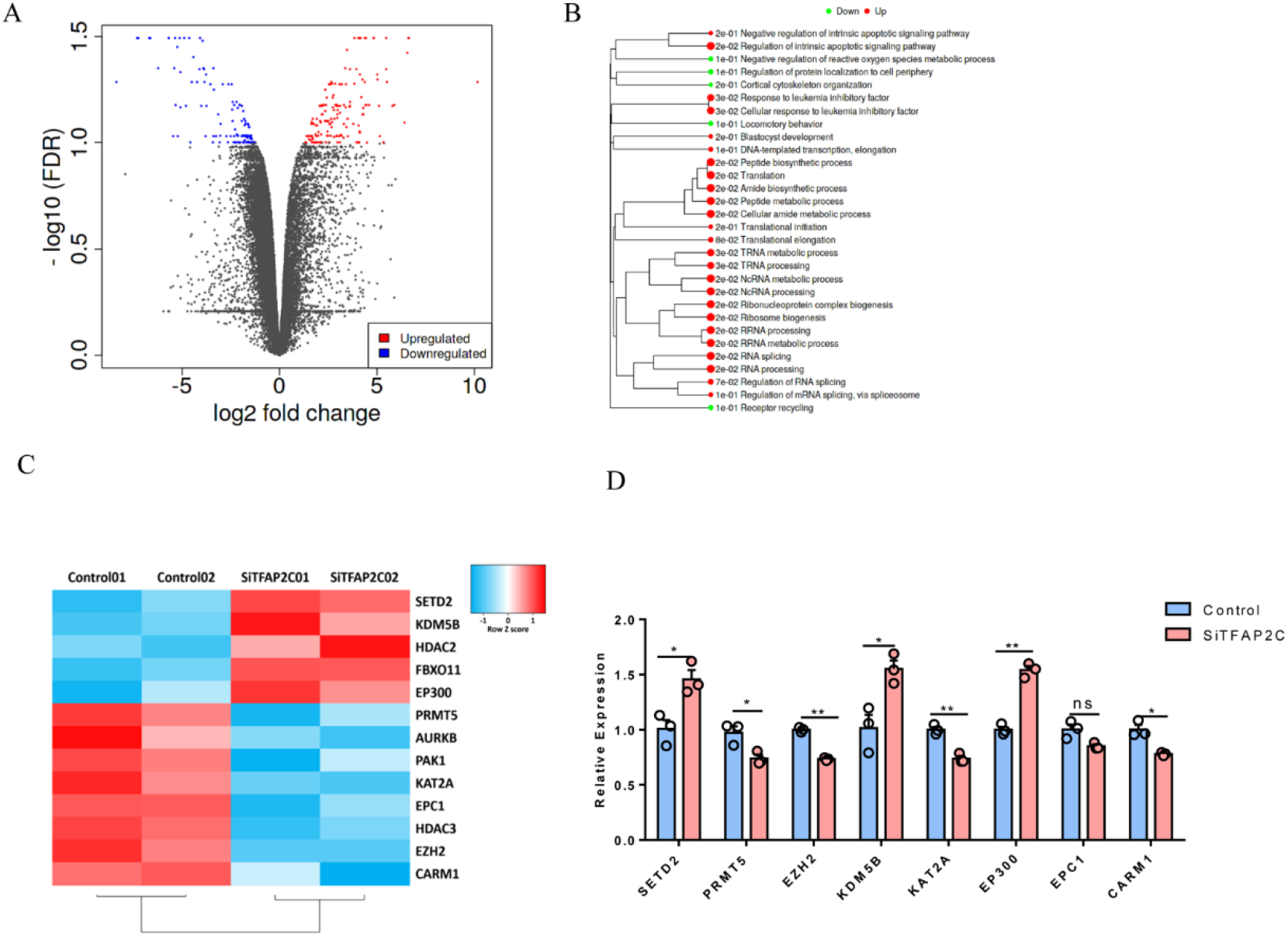
Interference with TFAP2C disrupted gene expression patterns. (A) The volcano showed differentially expressed genes between control group and *TFAP2C-KD* group, each dot represented a gene, the red dot indicated the upregulated gene and the blue dot indicated the downregulated gene, FDR<0.1, fold change>2. (B) Pathway enrichment of upregulated genes and downregulated genes. (C) Heatmap of main epigenetic genes in control group and *TFAP2C*-KD group. The color scale bar was shown at the right of the heatmap. Blue indicated the lowest expression, and red indicated the highest expression. (D) The significantly changed epigenetic genes, *P*\* <0.05, *P*\*\* <0.01. The experiment was independently repeated three times.

Next, we focused on the genes related to the epigenetic modification, as shown in figure 3C, the heatmap showed the expression pattern of primary epigenome genes. *SETD2* (*P*\*<0.05), *KDM5B* (*P*\*<0.05) and *EP300* (*P*\*\*<0.01) were increased significantly, and *EZH2* (*P*\*\*<0.01), *KAT2A* (*P*\*\*<0.01), *PRMT5* (*P*\*<0.05) and *CARM1* (*P*\*<0.05) were significantly decreased (figure 3D). These data revealed the knockdown of TFAP2C disturbed the transcription pattern during embryonic development and affected the expression of epigenetic modification genes.

### 2.4 SETD2 and KDM5B increased after *TFAP2C* knockdown

Next, we detected the expression of SETD2 and KDM5B by IF. As shown in figure 4A, the expression of SETD2 persisted at the 2-cell to 8-cell stage and decreased at the blastocyst stage. While SETD2 was significantly increased in 2-cell, 4-cell and 8-cell stages after interfering with *TFAP2C* (*P*\*\*<0.01). The KDM5B increased in the *TFAP2C-KD* group compared to the control group (*P*\*\*\*<0.001). These data indicated the knockdown of TFAP2C would induce the expression of SETD2 and KDM5B.

**Figure 4.**
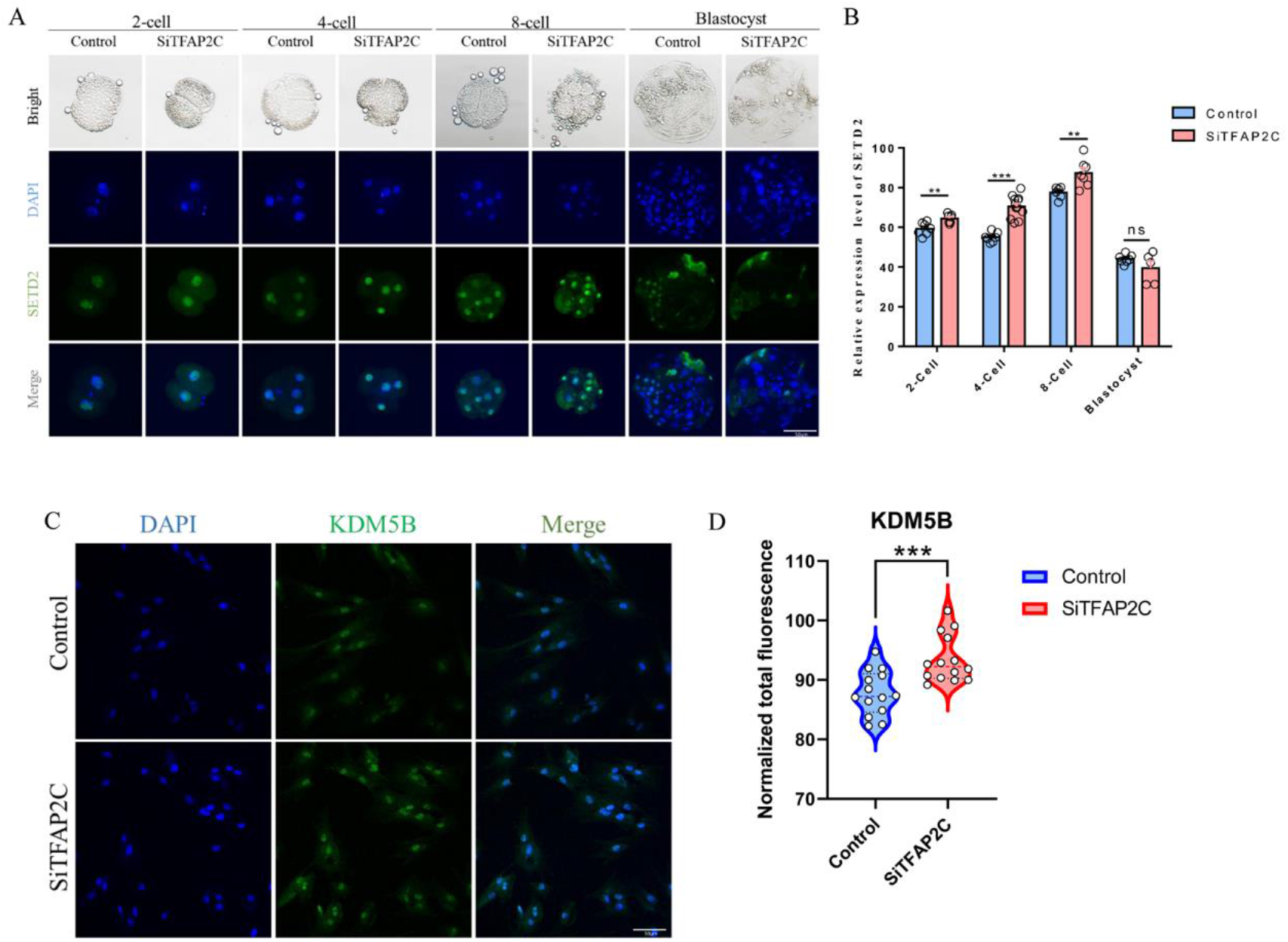
SETD2 and KDM5B increased after knocking down TFAP2C. (A) Immunofluorescence of SETD2 in 2-cell, 4-cell, 8-cell and blastocyst. The nuclei (blue) were stained with DAPI. The experiment was repeated three times independently with 6-10 embryos/stage /group /replicate. (B) Fluorescence intensity analysis of SETD2, *P*\*\* <0.01, *P* *** <0.001. (C) Immunofluorescence of KDM5B in fetal fibroblasts. The nuclei (blue) were stained with DAPI. The experiment was repeated three times independently. (D) Fluorescence intensity analysis of KDM5B, *P*\*\*\* <0.001.

### 2.5 The knockdown of TFAP2C induced abnormal epigenetic reprogramming in porcine embryos

Due to the abnormal expression of epigenetic genes in RNAseq, we detected the main histone modification level by IF. As shown in figure 5A and 5B, the IF staining results showed H3K4me3 was downregulated in the 2-cell (*P*\*<0.05) stage and 4-cell stage (*P\*\**<0.01) in the *TFAP2C-KD* group compared to the control group. H3K4me2 was downregulated in 2-cell (*P*\*<0.05) stage, 4-cell stage (*P\*\**<0.01) and blastocyst (*P\*\**<0.01) (figure 5C and figure 5D) in the *TFAP2C-KD* group. In addition, the H3K9me3 IF staining showed downregulated levels in the 4-cell stage (*P\*\**<0.01) and blastocyst in the *TFAP2C-KD* group (*P\*\**<0.01) (figure 6A and figure 6B). While the H3K36me3 was upregulated in the *TFAP2C-KD* group compared to the control group (figure 6C and figure 6D). Therefore, the knockdown of TFAP2C induced abnormal epigenetic modification.

**Figure 5.**
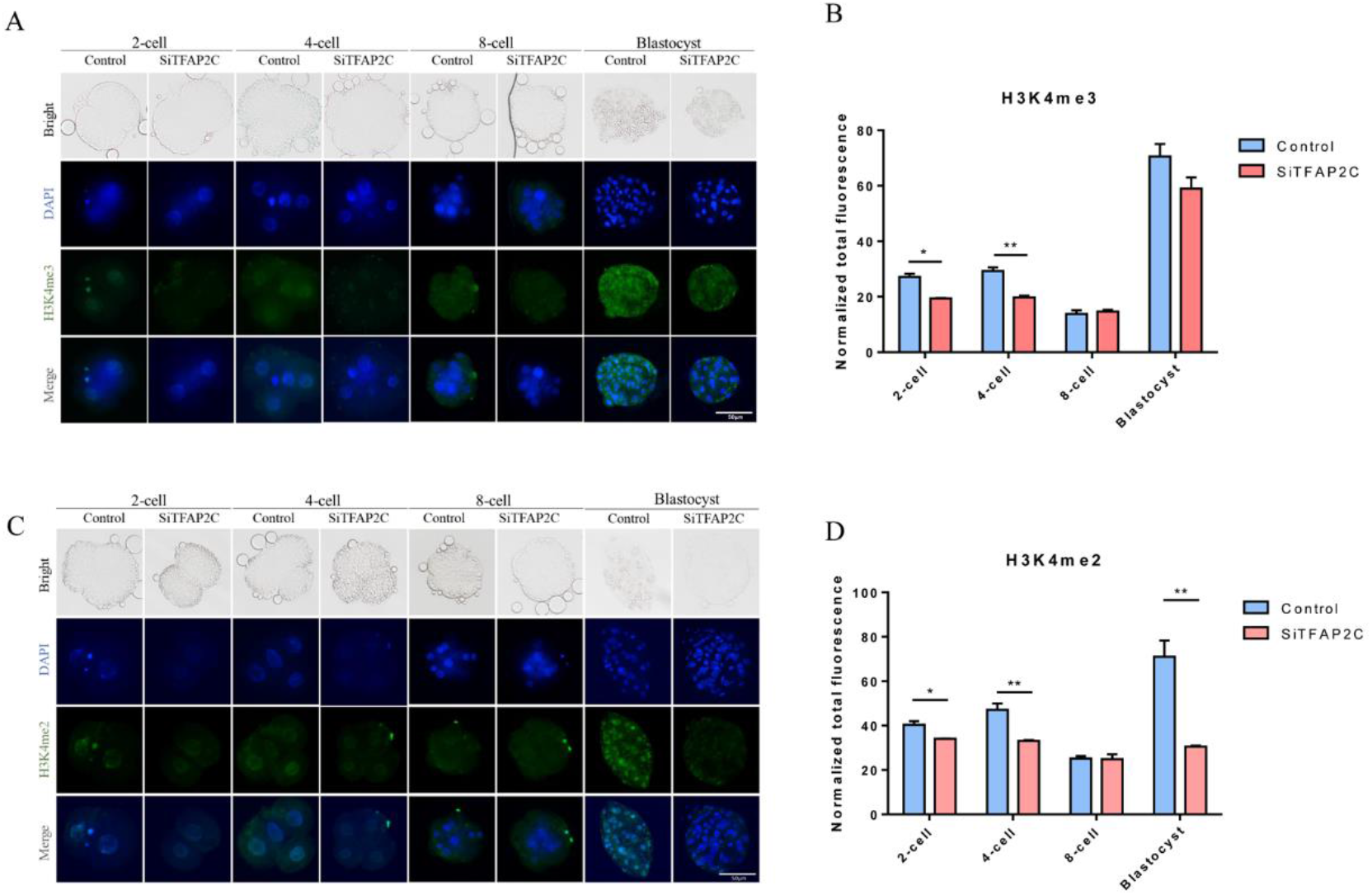
TFAP2C knockdown resulted in abnormal modifications of H3K4me3 and H3K4me2. (A) Immunofluorescence of H3K4me3 in 2-cell, 4-cell, 8-cell and blastocyst. The nuclei (blue) were stained with DAPI. The experiment was repeated three times independently with 6-10 embryos/stage /group /replicate. (B) Fluorescence intensity analysis of H3K4me3, *P*\* <0.05, *P*\*\* <0.01. (C) Immunofluorescence of H3K4me2 in 2-cell, 4-cell, 8-cell and blastocyst. The nuclei (blue) were stained with DAPI. The experiment was repeated three times independently with 6-10 embryos/stage /group /replicate. (D) Fluorescence intensity analysis of H3K4me2, *P*\* <0.05, *P*\*\* <0.01.

**Figure 6.**
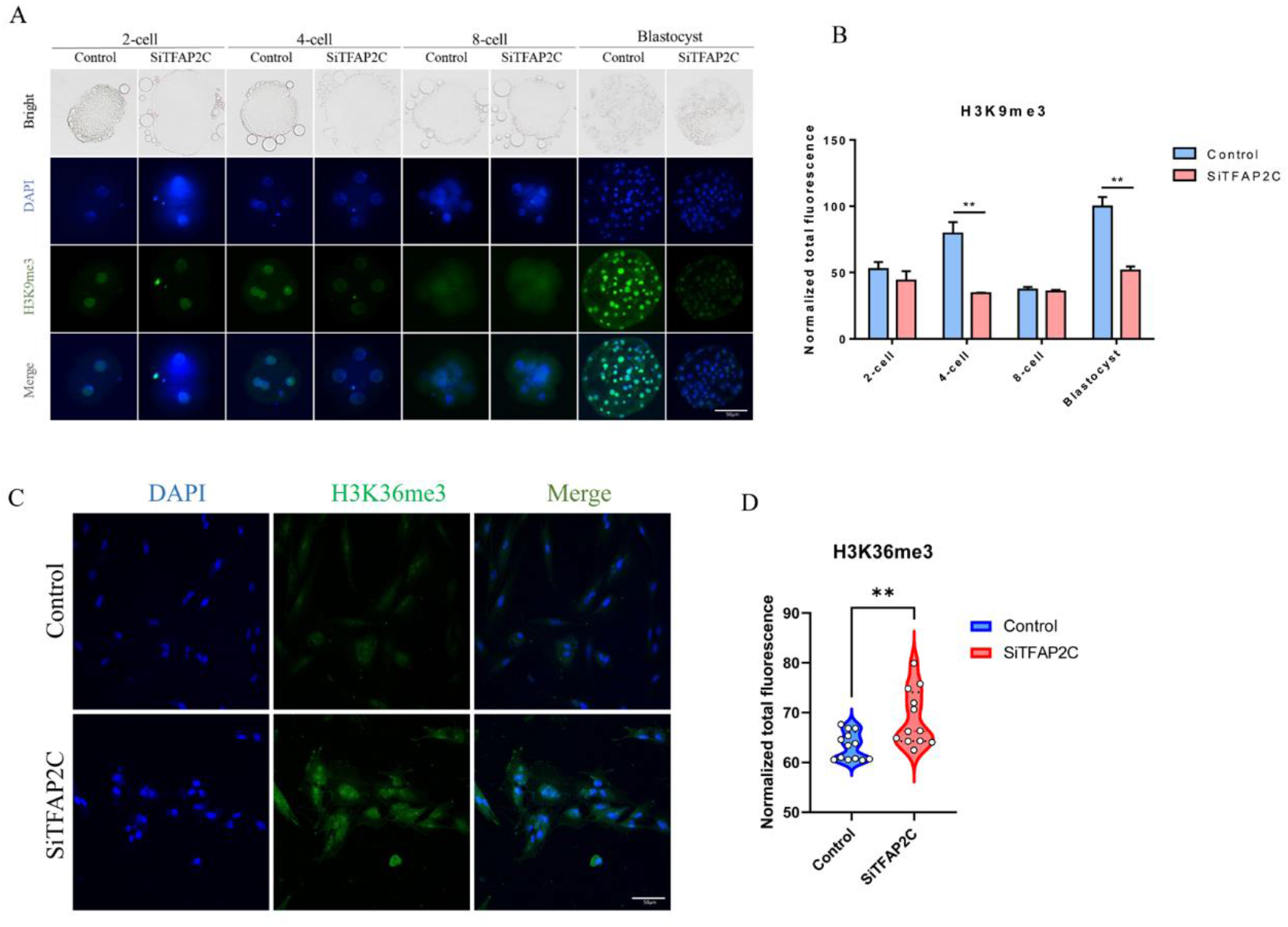
The knockdown of TFAP2C induced abnormal modifications of H3K9me3 and H3K36me3. (A) Immunofluorescence of H3K9me3 in 2-cell, 4-cell, 8-cell and blastocyst. The nuclei (blue) were stained with DAPI. The experiment was repeated three times independently with 6-10 embryos/stage /group /replicate. (B) Fluorescence intensity analysis of H3K9me3, *P*\*\*<0.01. (C) Immunofluorescence of H3K36me3 in fetal fibroblasts. The nuclei (blue) were stained with DAPI. The experiment was repeated three times independently. (D) Fluorescence intensity analysis of H3K36me3, *P*\*\*<0.01.

### 2.6 The knockdown of TFAP2C induced abnormal DNA methylation in porcine embryos

Next, we detected the 5mC/5hmC level in embryos, as shown in figure 7A, where IF staining showed the 5mC and 5hmC levels in 2-cell, 4-cell, 8-cell and blastocyst. The 5mC was upregulated in 2-cell (*P\*\**<0.01) and 4-cell stages (*P\*\**<0.01) (figure 7B), suggesting abnormal DNA methylation modification occurred in *TFAP2C*-KD embryos. Meanwhile, we detected the expression of DNMT1, as shown in figure 7C, the DNMT1 was mostly expressed in cytoplasm in the control group, while mostly expressed in nuclei in the *TFAP2C*-KD group. These data indicated the DNA methylation modification in *TFAP2C*-KD embryos was abnormal.

**Figure 7.**
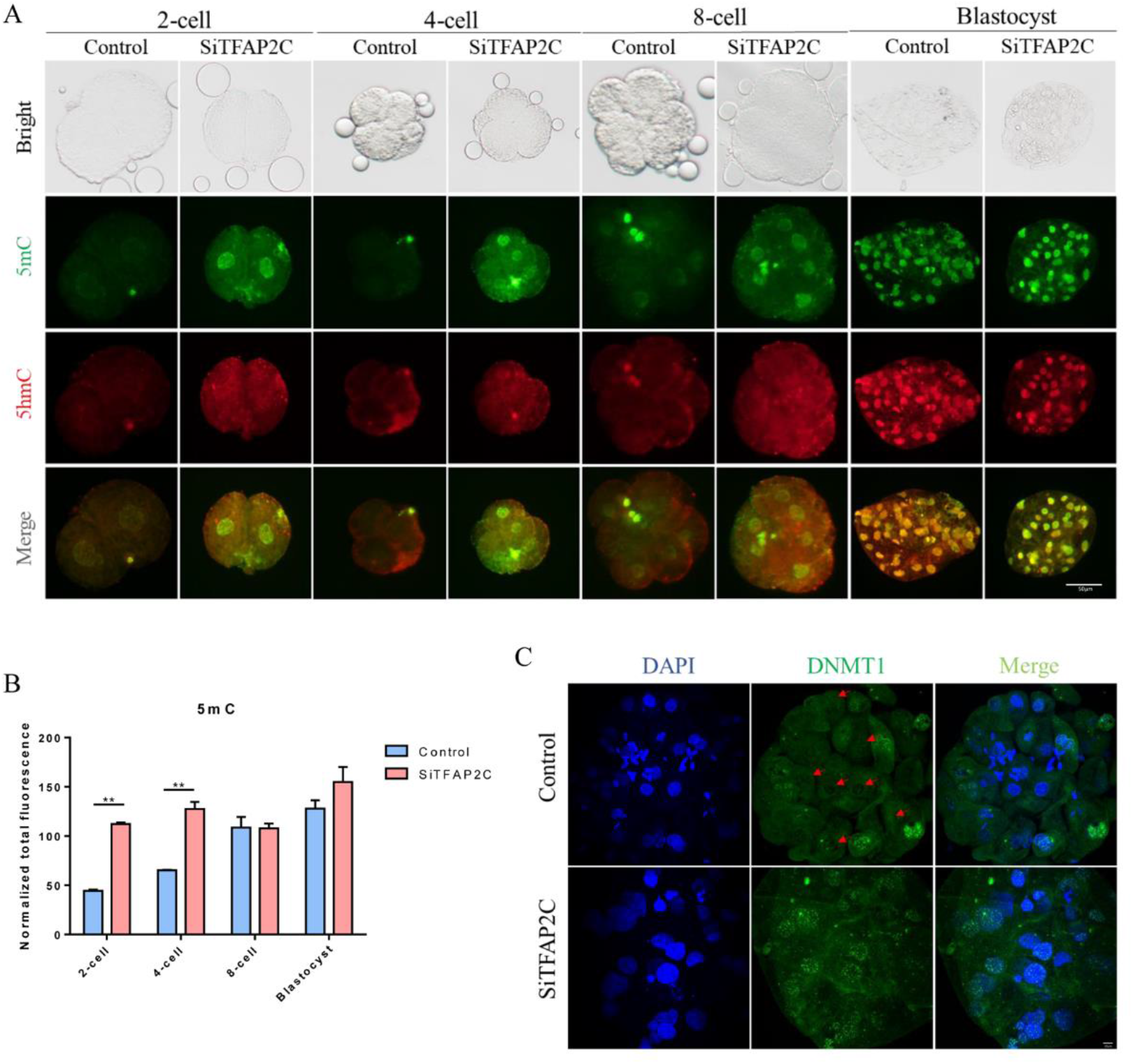
The knockdown of *TFAP2C* induced abnormal DNA methylation in embryos. (A) Immunofluorescence of 5mC/5hmC in 2-cell, 4-cell, 8-cell and blastocyst. The green indicated 5mC, and the red indicated 5hmC. (B) Fluorescence intensity analysis of 5mC, *P*\*\*<0.01. (C) Immunofluorescence of DNMT1 in blastocyst, the red arrow indicated the location of the nucleus. The experiment was repeated three times independently with 6-10 embryos/stage /group /replicate.

### 2.7 TFAP2C could bind to the promoter of *SETD2*

To figure out how TFAP2C regulates the expression of epigenetics genes, we predicted the promoter region binding sites of epigenetics genes based on the canonical TFAP2C-binding motif GCCNNNGGC (figure 8A). As shown in figure 8B, the TFAP2C-binding motif GCCNNNGGC exists in the promoter regions of many epigenetics genes, such as *SETD2, EP300, PRMT5, KAT2A, EPC1, EZH2, CARM1 etc*.. Next, we performed chromatin immunoprecipitation (ChIP)-PCR to confirm which promoter was occupied by TFAP2C. As shown in figure 8C, the promoter of *SETD2* was occupied with TFAP2C, indicating TFAP2C may regulate the expression of SETD2 by binding the promoter directly. Then, we mutated the classical pattern of TFAP2C in the *SETD2* promoter and results showed that the fluorescence activity of the WT group was significantly higher than that of the control group, while the fluorescence activity of the mutant group was higher than that of the WT group (figure 8D) (*P*\*\*<0.01), suggesting that TFAP2C may function as a blocking protein.

**Figure 8.**
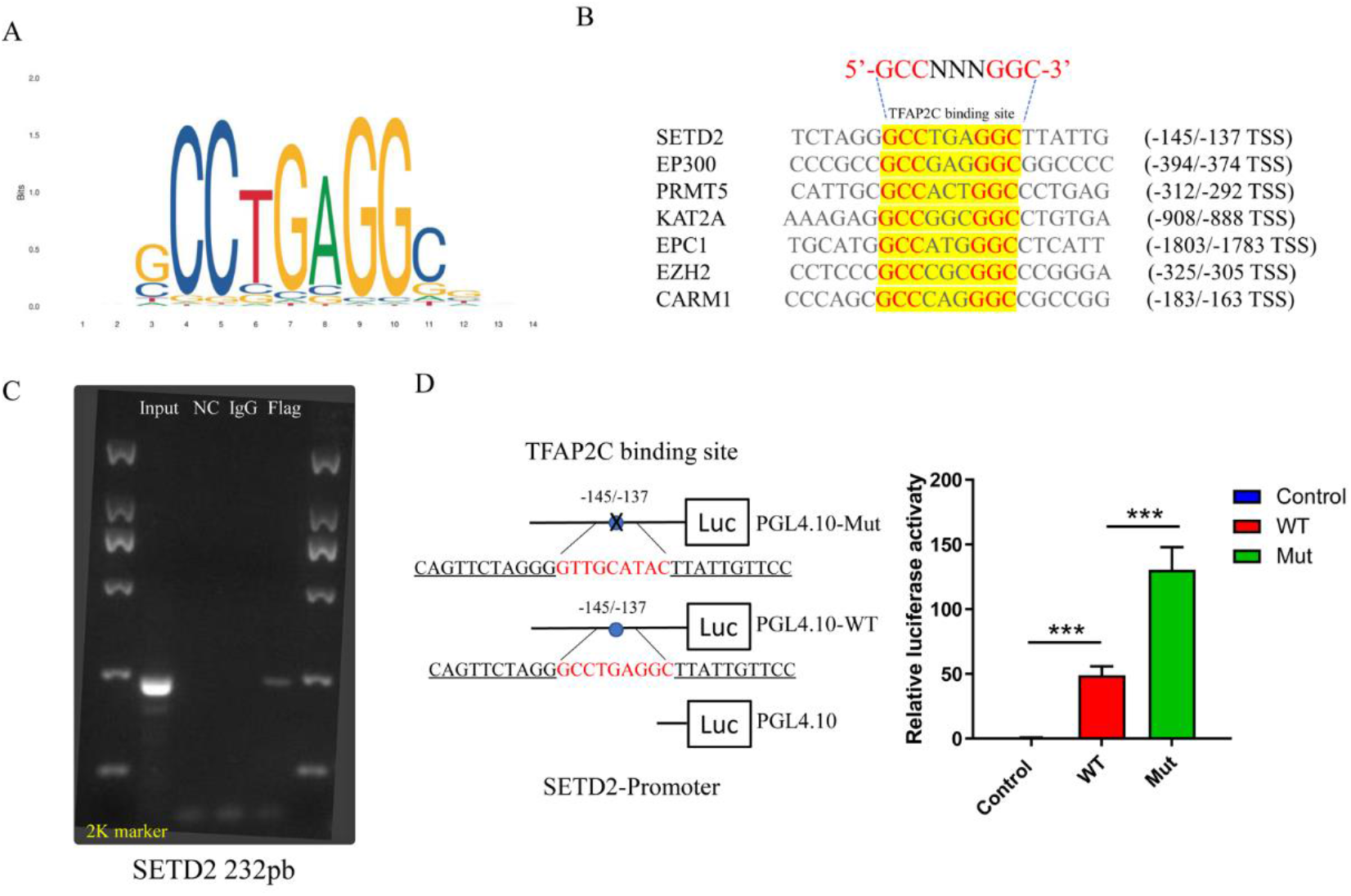
TFAP2C could bind to the promoter of *SETD2*. (A) The classical binding motifs of TFAP2C. (B) Prediction of promoter binding sites. (C) The result of ChIP assay, the lanes from left to right were Input, NC, IgG and Flag. (D) Dual luciferase reporter experiment.

### 2.8 High expression of TFAP2C had no significant effect on the expression of SETD2 and H3K36me3

To confirm the regulatory effect of TFAP2C on SETD2, we overexpressed TFAP2C and observed its effect on SETD2 and H3K36me3. As shown in figure 9A, we detected the expression of TFAP2C after injecting the TFAP2C mRNA into embryos, the results showed TFAP2C was increased in the mTFAP2C group (mRNA injection group) compared to the control group. We detected the protein expression by IF, which also increased (figure 9B). While there were no differences in SETD2 expression in embryos between the mTFAP2C group and the control group (figure 9C). When we detected the H3K36me3, it also had no significant differences (figure 9D). These data indicated that more expression of TFAP2C could not affect SETD2.

**Figure 9.**
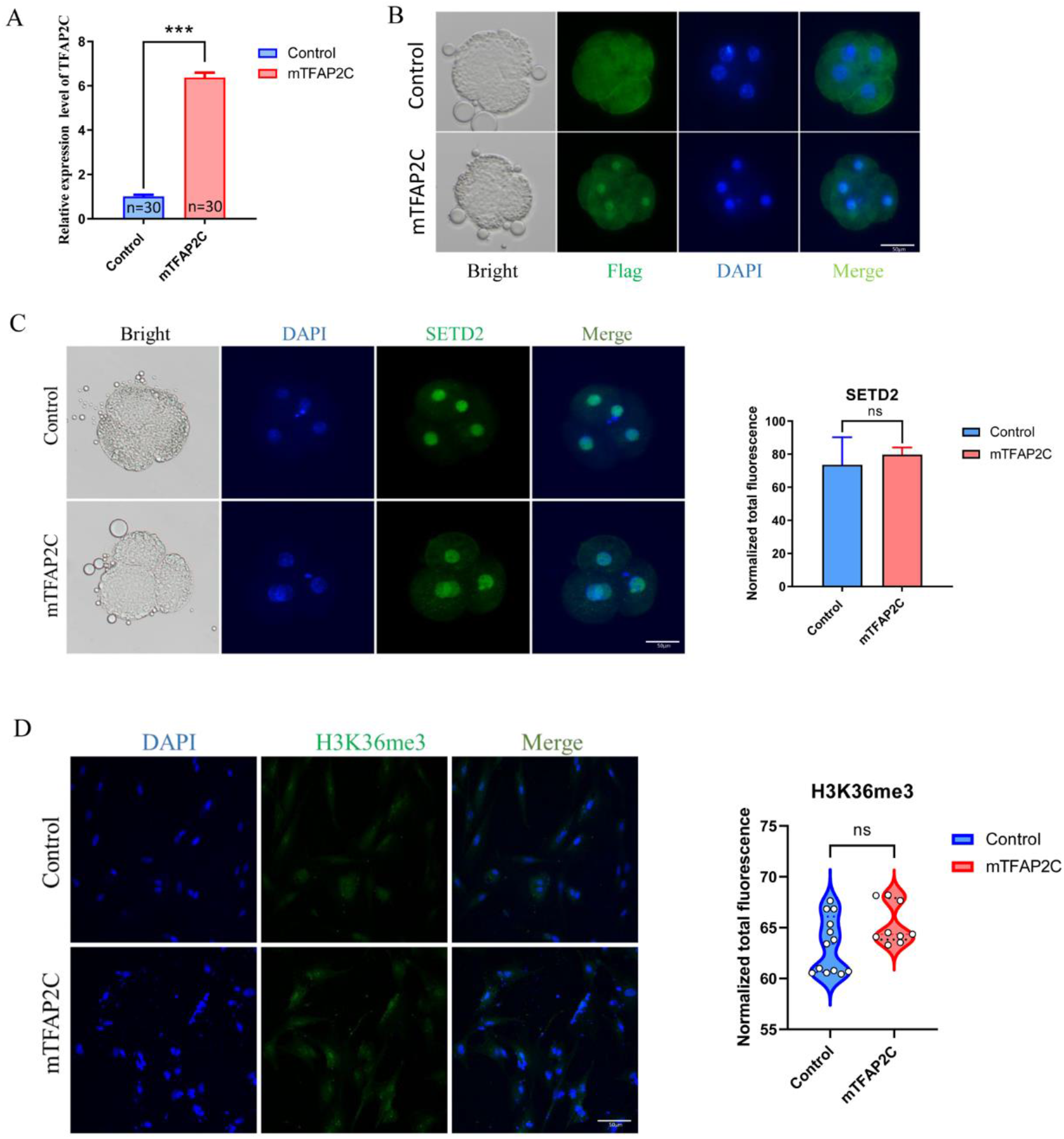
High expression of TFAP2C had no significant effect on the expression of SETD2 and H3K36me3. (A) qPCR analysis of *TFAP2C* expression of control and mTFAP2C groups at 48 h after mRNA injection (4-cell stage, n=30 per group). The experiment was independently repeated three times. (B) Immunofluorescence of Flag in 4-cell stage. The experiment was repeated three times independently with 6-10 embryos/group/replicate. (C) Immunofluorescence of SETD2 in 4-cell stage. The experiment was repeated three times independently with 6-10 embryos/group /replicate. Fluorescence intensity analysis was on the right of the image. (D) Immunofluorescence of H3K36me3 in fetal fibroblasts. The nuclei (blue) were stained with DAPI. The experiment was repeated three times independently. Fluorescence intensity analysis was on the right of the image.

## Discussion

Transcription factors are major developmental regulators that determine gene expression programs, and understanding their role is one of the central goals of developmental biology (20). Therefore, in this study, we systematically analyzed the expression patterns of major transcription factors during embryonic development using porcine embryos, and found that the timing of activation of most transcription factors highly overlapped with that of the zygotic genome activation (ZGA), so we hypothesized that this part of transcription factors might affect the embryonic development process by regulating gene expression.

TFAP2C has been shown to be associated with tumor progression (21–23), it is also a key factor in development. TFAP2C promotes somatic cell reprogramming by inhibiting c-Myc-dependent apoptosis and promoting mesenchymal-to-epithelial transition (24). Moreover, *Tcfap2c-induced* trophectoderm-like cells differentiate into trophectoderm derivatives in vitro and form trophoblast ectoderm in blastocyst in vivo, which is necessary for the establishment of an extra-embryonic trophectoderm maintenance program in mouse embryos (25). In post-implantation embryonic development, it has been reported that *TFAP2C* knockout mice were developmentally delayed at 7.5 days and did not survive beyond 8.5 days compared to controls (26). *TFAP2C* in mouse pre-implantation embryonic development declines sharply after fertilization and begins to rebound at the 4-cell stage (5). However, the authors did not investigate whether TFAP2C deficiency affects preimplantation embryo development. Aston *et al*. examined the expression of TFAP2C in bovine SCNT embryos and found that it was undetectable in oocytes and early embryos, while relatively high levels were detected in embryonic dysplasia and blastocysts (27). In contrast, in pig parthenogenetic activation (PA) embryos, interfering with *TFAP2C* effectively inhibited the development of PA embryos and blocked the embryos at the 8-cell stage (28). In our study, we found that *TFAP2C* expression was elevated starting at the 4-cell stage of porcine embryos and persisted until the blastocyst stage. Interfering with *TFAP2C* effectively inhibited the developmental efficiency of porcine embryos, and with significant differences at the 8-cell stage and induced apoptosis. The RNAseq results showed there were 1208 genes downregulated and 792 genes upregulated after TFAP2C knockdown, suggesting that the inhibited effect was due to the disrupted gene expression.

Histone modifications play a key role in the spatial and temporal regulation of mammalian gene expression(29). Shao *et al*. reported that the increased expression of H3K4me2 at 2-cell stage caused abnormal activation of embryonic gene expression and further reduction of developmental efficiency in mice(30). In porcine embryos, H3K4me3 and H3K27me3 decrease steadily from zygotes to the lowest levels in morula and subsequently increase again in blastocysts(31). The abnormal modification of H3K4me3 was correlated with abnormal gene expression in extraembryonic tissues(32). Histidine-lysine N-methyltransferase SETD2 is an H3K36me3 methyltransferase(33), Xie *et al*. reported that H3K36me3 could recruit KDM5B to gene bodies to promote demethylation of H3K4me3 (34). However, no studies have reported the regulatory role of *TFAP2C* in embryos on epitope-modifying enzymes. Our results showed that interference with *TFAP2C* affected gene expression of epigenetic modifying enzymes, with elevated expression of *KDM5B, SETD2* and *EP300* and decreased expression of *PRMT5*, *CARM1*, *EZH2*, *KAT2A* and *ECP1*. Meanwhile, the modification levels of H3K4me3 and H3K4me2 were significantly decreased and the modification level of H3K36me3 was significantly increased at the 2-cell and 4-cell stages in the *TFAP2C-KD* group, suggesting that the lack of *TFAP2C* caused abnormal histone modifications, resulting in impaired embryonic development.

DNA methylation is also a type of epigenetic modification, which is involved in various biological processes in mammals, including the silencing of transposable factors, regulation of gene expression, genomic imprinting, and X chromosome inactivation. It has been shown that DNA methylation and histone modification interacted with each other, the methylation of CpG islands related to the methylation status of H3K4 (11), the levels of methylated H3K4 (H3K4me3) tend to be inversely correlated with DNA methylation (19). However, studies on TFAP2C-mediated DNA methylation changes have not been reported. Our results showed increased levels of 5mC modifications in the 2-cell and 4-cell stages of IVF embryos after interference with *TFAP2C*. And the modification level of H3K4me3 was decreased. In addition, DNMT1 appeared to be nucleated in blastocysts lacking *TFAP2C*. It indicated that TFAP2C could affect DNA methylation level, which in turn affects embryonic development.

*TFAP2C* has been shown to regulate the pluripotency regulator OCT4 by binding to enhancers that are essential in hESCs (35). *TFAP2C* is induced in mouse fibroblasts during iPSC production and acts as a promoter of iPSC formation, regulating its expression by binding to the promoter region of Cdh1(7). It has also been shown that a ternary complex of TFAP2C, oncoprotein Myc, and histone H3 (H3K4me3) demethylase KDM5B exists at the proximal promoter, thereby regulating CDKN1A expression to control the cell cycle (36). In pig embryos, Lee *et al*. reported that OCT4 transcript levels were elevated in pig *TFAP2C*-KD 8-cell embryos and that *TFAP2C* may be involved in regulating OCT4 (37), however, the authors did not perform further experiments to verify this regulatory relationship. In this study, we reported for the first time that TFAP2C could bind to the promoter region of *SETD2* to regulate its expression, and the two showed an inverse correlation. *SETD2*-mediated levels of H3K36me3 were negatively correlated with levels of *EZH2*-catalyzed H3K27me3, suggesting that SETD2 can methylate *EZH2* and promote EZH2 degradation (38). Meanwhile, SETD2-catalyzed H3K36me3 modification is involved in crosstalk with other chromatin markers, including antagonism of H3K4me3 and H3K27me3, leading to de novo DNA methylation through recruitment of DNA methyltransferases 3A and 3B (33, 39). Therefore, we speculated that TFAP2C interference in our results caused elevated SETD2 expression, which mediated increased H3K36me3 modification.

Simultaneously increased SETD2 caused degradation of EZH2 on the one hand and recruited KDM5B for H3K4me3 demethylation on the other. Although previous study reported that the knockdown of SETD2 inhibited porcine embryonic development (40), our result showed the high expression of SETD2 still not conducive to embryonic development, suggesting the abnormal expression of SETD2 was detrimental to embryonic development, and only appropriate SETD2 gene expression level was beneficial to normal embryonic development.

### Conclusion

Our results suggest that TFAP2C is a key TFs regulating early embryo development in pigs. It can affect embryonic development by regulating epigenetic modifying enzymes such as KDM5B, SETD2 *ect*. and epigenetic modifications such as H3K4me3, H3K36me3 and DNA methylation modifications *ect*.. In addition, we report for the first time that the classical binding motif of TFAP2C can bind to the *SETD2* promoter region and regulate its expression, thus affecting embryonic developmental processes. These findings provide a theoretical basis for improving developmental efficiency and animal production.

## Materials and methods

### 4.1 TFs isolation and analysis

The RNAseq data of IVV embryos we used is the data we have uploaded to NCBI previously (https://submit.ncbi.nlm.nih.gov/subs/sra/SUB10711406/PRJNA 783716), under accession IDs: MII_1 (SRR17041081), MII_2 (SRR17041080), MII_3 (SRR17041074), IVV_2C_1 (SRR17041073), IVV_2C_2 (SRR17041072), IVV_2C_3 (SRR17041071), IVV_4C_1 (SRR17041070), IVV_4C _2 (SRR17041069), IVV_4C_3 (SRR17041068), IVV_8C_1 (SRR17041067), IVV _8C_2 (SRR17041079), IVV_8C_3 (SRR17041078), IVV_Bla_1 (SRR17041077), IVV_Bla_2 (SRR17041076), IVV_Bla_3 (SRR17041075). Transcription factors were extracted according to the genes in table S1. The heatmap was constructed by TBtools.

### 4.2 Antibodies and Chemicals

H3K4me2 (Abcam; ab7766), H3K4me3 (Abcam; ab8580), and H3K9me3 (Abcam; ab8898), H3K36me3 (Bioss; bs-3768R), SETD2 (Affinity, AF7552), KDM5B (omnimabs; OM260516), 5mC (Eurogentec; BI-MECY-0100), 5hmC (Active motif; 39769), DNMT1 (Invitrogen; MA5-16169), Alexa Fluor 488 goat anti-rabbit (Invitrogen; A-11008) and/or Alexa Fluor 488/594 goat anti-mouse (Invitrogen; A32723/A-11020) were used in this study. All the chemicals used to culture were purchased from Sigma-Aldrich (St. Louis, MO, USA) unless otherwise stated.

### 4.3 The collection and in vitro maturation of porcine oocytes

The pig ovaries we used were taken from the same local slaughterhouse (Changchun Huazheng, Jilin, China), and oocytes were transported to the laboratory within 2 hours and were kept in 0.9% NaCl supplemented with 200 IU/mL penicillin and streptomycin at 35-36.5 °C.

The follicular fluid containing cumulus-oocyte complexes (COCs) from 3-6 mm ovarian follicles were aspirated using an 18-gauge needle. COCs with at least three layers of cumulus cells were selected, washed three times in manipulation fluid (TCM-199 supplemented with 0.1% polyvinyl alcohol), and then cultured in in vitro maturation (IVM) media. Approximately 200 COCs were each cultured in a 1 mL drop of maturation medium (TCM-199 supplemented with 10 μg/mL epidermal growth factor, 0.5 μg/mL porcine luteinizing hormone, 0.5 μg/mL porcine follicle-stimulating hormone, 26 mM sodium bicarbonate, 3.05 mM glucose, 0.91 mM sodium pyruvate, 0.57 mM cysteine, 0.1% PVA, 10% foetal calf serum, 75 mg/mL penicillin G and 50 mg/mL streptomycin) for 22-24 h and then cultured with hormone-free maturation medium (The formula is consistent with the previous maturation medium without 10 μg/mL epidermal growth factor, 0.5 μg/mL porcine luteinizing hormone and 0.5 μg/mL porcine follicle-stimulating hormone) for 20 h at 38.5 °C and 5% CO2. Then, cumulus cells were removed from oocytes with manipulation fluid supplemented with 0.2% hyaluronidase. The oocytes with polar body 1 (PB1) were considered matured and used for the following experiments.

### 4.4 Microinjection/ Cytoplasmic injection

SiRNA was injected by the microinjection meter (Eppendorf, FemtoJet 4i, USA). 5-10 pL of 25 nM *TFAP2C* siRNA or NC siRNA were injected into MII oocytes, and then they were incubated with semens in PGM (porcine gamete medium: 100 ml water with 0.6313 g NaCl, 125 0.07456 g KCl, 0.00477 g KH2PO4, 0.00987 g MgSO4·7H2O, 0.2106 g NaHCO3, 0.07707 g 126 CaC6H10O6·5H2O, 0.0187 g D-Glucose, 0.3 g PVA, 0.00242 g Cysteine, 0.04504 g C7H8N4O2, 0.0022 127 g C3H3NaO3, and 100 μl/mL penicillin/streptomycin). The *TFAP2C* siRNA and NC siRNA were designed and synthesized in Sangon biotech (shanghai, China). The sequences were as followed:

*TFAP2C* siRNA1: sense: GCCCUGAUAGUCAUAGAUATT

anti-sense: UAUCUAUGACUAUCAGGGCTT;

*TFAP2C* siRNA2: sense: CCGCACAGCAAGUGUGUAATT

anti-sense: UUACACACUUGCUGUGCGGTT

NC: sense: UUCUCCGAACGUGUCACGUTT

anti-sense: ACGUGACACGUUCGGAGAATT

### 4.5 In vitro fertilization of oocytes

Fresh semen was collected from the Jilin University pig farm. Density gradient centrifugation was used to wash them. In brief, percoll was configurated in the concentration of 90% and 45%, 2 ml semen was added and centrifuged at 300 g for 20 minutes, the supernatant was removed. Then 4 ml DPBS was added and centrifuged at 300 g for 10 min. The sperm were resuspended with PGM. Sixty denuded matured oocytes were cultured in 400 μl PGM with a final sperm concentration of 1.6×10^5^ - 5.0×10^5^ sperm/ml, at 38.5 °C and 5% CO_2_ for 5-6 h. After washing off the adherent sperm, the fertilized oocytes were transferred to PZM-3.

### 4.6 RNA-seq

Smart-Seq2 method was used to amplify each sample (6-8 embryos for each group) according to the manufacturer’s instructions. RNA concentration of library was measured by Qubit 2.0 Fluorometer (Life Technologies, CA, USA). Agilent Bioanalyzer 2100 system was used to assess the insert size and the quality of the amplified products was evaluated according to the detection results. Amplified product cDNA was used as input for the library construction of transcriptome. After the library construction, Agilent Bioanalyzer 2100 system was used to assess insertion size, and Taq-Man fluorescence probe of an AB Step One Plus Real-Time PCR system (Library valid concentration > 10 nM) was used to quantify the accurate insertion size. Clustering of the index-coded samples was performed using a cBot cluster generation system and the HiSeq PE Cluster Kit v4-cBot-HS (Illumina). Then, the libraries were sequenced by Zhejiang Annoroad Biotechnology (Beijing, China) on an Illumina platform, and 150 bp paired-end reads were generated. We used STAR to compare transcriptome data and cuffLinks for quantitative splicing. DEmRNAs and DElncRNAs were identified by the IDEP website (http://bioinformatics.sdstate.edu/idep/). Genes with false discovery rates (FDRs) ≤ 0.1, |log2FC| ≥ 1.5 and P-value ≤ 0.01 were selected as candidate genes. The data was showed in table S2.

### 4.7 RNA isolation and quantitative PCR

Total RNA was extracted and complementary DNA (cDNA) was synthesized with SuperScriptTM IV CellsDirect™ cDNA Synthesis kit (11750350, Invitrogen, USA) following the manufacturer’s instructions. qPCR was performed with FastStart Essential DNA Green Master (06924204001, Roche, USA) via a StepOnePlus Real-Time PCR system. Primer sequences were listed in Table S3.

The results were analyzed using the 2^-ΔΔCT^ method. The qPCRs were all repeated three times.

### 4.8 Immunofluorescence (IF) staining

Embryos were washed with PBS containing 0.1% polyvinylpyrrolidone. The zona pellucida (ZP) of embryos was dissolved in acidic Tyrode solution (pH 2.5). After washing in PBS-PVP, embryos were fixed with 4% paraformaldehyde for 30 min in the dark. After being washed in PBS-PVP, embryos were permeabilized with 0.2% Triton X-100/PBS (v/v) for 20 min and then blocked with 2% BSA/PBS for 1 h. For 5mC/5hmC staining, the embryos were treated with 4N-HCl and Tris-HCl for 30 minutes each before BSA blocking. Embryos were incubated with primary antibodies overnight at 4 °C. After being washed in PBS-PVP, embryos were stained with second antibody at 37 °C for 2 h in dark. Then DNA was stained with 10 μg/mL DAPI for 15 min. All samples were observed under a Nikon Eclipse Ti-U microscope equipped with appropriate filters (Nikon, Tokyo, Japan) after mounting. Colour images were captured using a DS-Ri2 CCD camera (Nikon, Tokyo, Japan) and analysis software (NIS-Elements BR; Nikon, Tokyo, Japan). The same exposure times and microscope settings were used for all captured images.

Evaluation of fluorescence intensity of individual images was performed using ImageJ software (National Institutes of Health, Bethesda, MD). The cytoplasmic background fluorescence intensity was measured as an average intensity level within the cytoplasmic area. Thereafter, the correction for the cytoplasmic background was carried out and the background-subtracted images were used for further analysis (41). All operations were carried out at room temperature unless otherwise specified.

### 4.9 Apoptosis assays on the blastocysts

Blastocysts were collected on the seventh day of incubation, the zona pellucida was removed by treatment with acidic Tyrode solution (pH 2.5), and embryos were fixed in 4% paraformaldehyde for 30 min. Fixed blastocysts were permeabilized for 30 min using 0.2% Triton X-100, washed three times with PBS-PVP, and incubated at 37 °C in the dark for 1 h with terminal deoxynucleotidyl transferase dUTP nick end labeling (TUNEL) solution from the In Situ Cell Death Detection Kit (Roche, Mannheim, Germany). The embryos were washed three times with PBS-PVP, and incubated for 10 min with DAPI to stain the nuclei. The stained embryos were mounted between a cover slip and a glass slide and examined under a fluorescence microscope.

### 4.10 Western blotting

Total protein was extracted from 80-100 embryos (4-cell stage) using RIPA with proteasome inhibitor, mixed with 4x SDS loading buffer and incubated in boiling water for 5 minutes. Equal amounts of protein were resolved by SDS-PAGE, the antibodies were as follows: TFAP2C (Sigma-Aldrich, AV38284), Vinculin (Proteintech, 26529-1-AP).

### 4.11 Transfection of porcine fetal fibroblasts

Fibroblasts were seeded into 6-well plates the day before transfection so that the confluence of cell growth was 30-40%. Cells were washed 2 times with pre-warmed PBS before transfection and 1 ml of medium without double antibodies was added and placed in the incubator. We used Promage’s plasmid transfection reagent, using 4:1, 3:1, and 2:1 ratios respectively according to the instructions for pre-fixing, and in our transfection experiments we used a ratio of 2:1. The transfection reagent, which had been incubated for 15 min each at room temperature, was mixed with the opti-MEM mixture, and the plasmid was mixed with the opti-MEM mixture and continued to incubate at room temperature for 10 min. After completion, the mixture was added drop-wise to the Petri dishes and the samples were collected after 48 h for testing.

### 4.12 ChIP

ChIP experiments were performed according to the instructions (17-295, Millipore). We set four groups as Input, NC, IgG, Flag (Fusion protein of TFAP2C and Flag), and finally, the DNA samples were amplified by PCR. The primers for PCR were listed in table S4.

### 4.13 Dual luciferase assay

The dual fluorescein assay was performed by the Promega kit, following the instructions for the experiment, and the firefly and sea kidney fluorescence values were measured by enzyme marker, and the firefly fluorescence value/sea kidney fluorescence value was the relative fluorescence intensity.

### 4.14 Statistical analysis

Data were presented as the means ± SEMs. The experiments were repeated at least three times in triplicate. Statistical analysis was performed using GraphPad Prism 9.0.0 (GraphPad Software, U.S.A.). *P*\*<0.05, *P*\*\*<0.01 and *P*\*\*\*<0.001 was considered statistically significant.

## Abbreviations

TFs: Transcription factors
KD: Knock down
ESCs: Embryonic stem cells
TSS: Transcription start site
SAM: S-adenyl methionine
TET: Ten-eleven translocation enzymes
IVV: In vivo fertilization
IVF: In vitro fertilization
SCNT: Somatic cell nuclear transfer
COCs: Cumulus-oocyte complexes
IVM: In vitro maturation
PB1: Polar body 1
FDRs: False discovery rates
FC: Fold change
ZGA: Zygotic genome activation
IF: Immunofluorescence
ZP: Zona pellucida

## Data Availability Statement

The data that support the findings of this study are available in the materials and methods and supplementary materials of this article.

## Bioethics

The experimental protocol of this study was approved by the Animal Care and Use Committee of the First hospital of Jilin University (2019-099).

## Authors’ contributions

Ziyi Li, Daoyu Zhang and Sheng Zhang conceived and designed the research; Daoyu Zhang, Di Wu, and Yongfeng Zhou performed the research and acquired the data; Daoyu Zhang, Meng Zhang analyzed and interpreted the data; Ziyi Li, Daoyu Zhang and Di Wu wrote the manuscript. All authors were involved in drafting and revising the manuscript.

## Supporting information

**S1 File: List of transcription factors.**

**S2 File: The RNAseq data.**

**S3 File: The primers used for qPCR.**

**S4 File: The primers used for ChIP PCR.**

